# Effects of mechanical loading on cortical defect repair using a novel mechanobiological model of bone healing

**DOI:** 10.1101/190819

**Authors:** Chao Liu, Robert Carrera, Vittoria Flamini, Lena Kenny, Pamela Cabahug-Zuckerman, George Benson, Daniel Hunter, Bo Liu, Gurpreet Singh, Philipp Leucht, Kenneth A. Mann, Jill A. Helms, Alesha B. Castillo

## Abstract

Mechanical loading is an important aspect of post-surgical care. The timing of load application relative to the injury event is thought to differentially regulate repair depending on the stage of healing. Here, we show using a novel mechanobiological model of cortical defect repair that daily loading (5 N peak load, 2 Hz, 60 cycles, 4 consecutive days) during hematoma consolidation and inflammation disrupts the injury site and activates cartilage formation on the periosteal surface adjacent to the defect. We also show that daily loading during the matrix deposition phase enhances both bone and cartilage formation at the defect site, while loading during the remodeling phase results in an enlarged woven bone regenerate. All loading regimens resulted in abundant cellular proliferation within the regenerate and at the periosteal surface and fibrous tissue formation directly above the defect. Stress was concentrated at the edges of the defect during exogenous loading, and finite element (FE)-modeled longitudinal strain (ε_*zz*_) values along the anterior and posterior borders of the defect (~2200 με) were an order of magnitude larger than strain values on the proximal and distal borders (~50-100 με). These findings demonstrate that all phases of cortical defect healing are sensitive to physical stimulation. In addition, the proposed novel mechanobiological model offers several advantages including its technical simplicity and its well-characterized and spatially confined repair program, making effects of physical and biological interventions more easily assessed.

## 1.0 Introduction

Mechanical loading regulates skeletal adaptation and fracture repair [1–3], and early weight-bearing is an important factor in orthopaedic post-surgical care [4]. Compressive strain, tensile strain, hydrostatic pressure, shear strain and fluid flow are all important modes of mechanical stimulation [5–9], though the rules governing the relationship between the timing of load application relative to the stage of healing, and underlying cell and molecular signaling, are poorly understood. Previous studies using osteotomy and segmental defect models showed that application of exogenous loading immediately after injury suppressed bone formation [10–12]. When loading was applied after a delay of days to weeks, both cartilage and bone formation increased relative to non-loaded controls [10–13]. In these models, the presence of fixation hardware (e.g., intramedullary pin, plates, external fixator) can exert secondary effects on healing, which may mask direct effects of exogenous mechanical loading.

To address this limitation, we utilized a self-stabilizing bone repair model that undergoes a well-defined bone repair program and is amenable to precisely controlled loading regimens at defined stages of healing. Here we describe this model, which couples a stable cortical defect [14,15] with a precisely-controlled mechanical stimulus. We present outcomes from loading at early, intermediate, and from late stages of repair. The data shown here support our hypothesis that loading during the early stages of repair suppress osteogenesis and enhance cartilage formation, while loading during the intermediate and later stages of repair enhance both cartilage and bone formation. In addition, we show that cellular proliferation in mechanically loaded defects is abundant within the regenerate and at the periosteal surface, suggesting that loading enhances both proliferation and differentiation of osteochondroprogenitors in the context of healing.

## 2.0 Materials and Methods

### 2.1 Experimental design

The NYU Institutional Animal Care and Use Committees approved all procedures. Sixteen-week-old C57BL/6 female mice were obtained from The Jackson Laboratory (Bar Harbor, ME) and had access *ad libitum* to standard mouse chow and water for the duration of the study. All mice underwent bilateral monocortical tibial defect surgery. Three mice were used for generation of *ex vivo* load-strain calibration curves and development of linear elastic finite element (FE) models of whole tibiae for estimating strain components at the defect site. The remaining mice were subjected to *in vivo* tibial axial compressive loading for four consecutive days beginning on post-surgical day (PSD) 2, 5, or 10. Non-loaded defects served as controls. All mice underwent *in vivo* longitudinal micro-computed tomographic (microCT) scanning of bilateral defects on PSD 2, 5, 8, 10 and 14 or until euthanized (see *In vivo longitudinal microCT*). Groups of mice were euthanized at PSD 2, 5, 10 or 14 for histological analyses.

### 2.2 Monocortical defect surgery

Under isoflurane anesthesia, a small skin incision was made over the midline of the anteromedial aspect of the tibia. A 1.0 mm circular defect centered between the tibiofibular junction (TFJ) and the tibial tubercule (Figure 1a) was made using a precision surgical drill. The defect was located 4.3 mm below the proximal articulating surface of the proximal tibia. After saline irrigation, the incision was closed with 7-0 nylon sutures. Mice were transferred into a clean cage positioned on a 37°C heating pad. Buprenorphine was administered immediately before surgery and at 6, 24, 48, and 72 hours after surgery.

**Figure 1.**
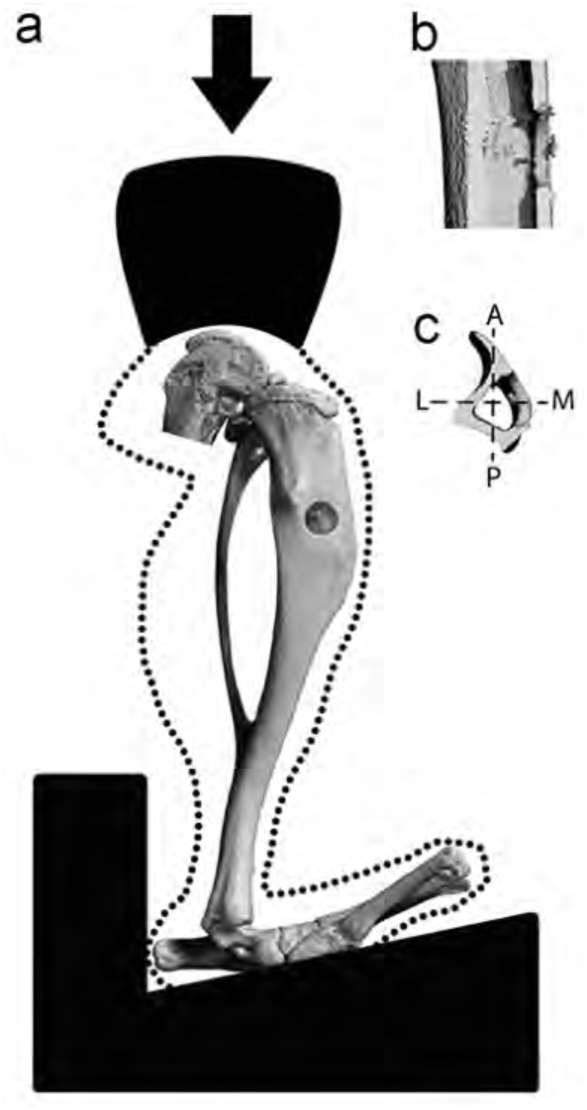
Axial compressive cyclic loading of a mouse tibia containing a monocortical defect. (a) Anteromedial, (b) sagittal, and (c) transverse views of the defect. Axes in the transverse view are A (anterior) – P (posterior) and L (lateral) – M (medial).

### 2.3 Finite element analysis with model validation

Linear elastic FE models of three individual tibiae at PSD 2, each containing a 1.0 mm cortical defect, were developed as representative experimental cases. Following euthanasia, the tibial surfaces were exposed and single axis strain gages (EA-06-015DJ120, Micro-Measurements, Wendell, NC) were applied 1 mm proximal and distal from the defect, with orientation coincident with the long axis of the tibia (Figure 2a). Limbs were loaded in axial compression to peak loads of 1 to 10 N in 1 N increments, and the longitudinal strains were recorded. Prior to microCT scanning, aluminum foil inserts were placed at the center of the knee loading cup and on the hind paw loading fixture. MicroCT scans (Scanco vivaCT 40) of the limbs were used to create 3D solid models of the tibia in MIMICS (Materialise, Plymouth, MI). Scan sets were oriented in MIMICS such that the centroids of the proximal and distal foil markers were aligned in the longitudinal direction. In this way, the orientation of the scan set was consistent with the axial orientation in the loading fixture. Voxel-based finite element meshes were created (432,000 elements, 530,000 nodes) with 0.03mm edge length. Homogenous and isotropic material properties were assigned to the tibia with a 17 GPa elastic modulus and 0.3 Poisson’s ratio. Linear elastic finite element analysis was performed in MSC Marc (MSC Software Co, Newport Beach, CA). Axial compressive loads were applied to small contact patches on the medial and lateral plateau regions proximally. Distally, the tibia was fixed to prevent translation in x, y, and z directions with nodal ties applied to distal contact regions (Figure 2a). In addition, translations in the x and y directions were prevented at a small segment of the proximal tibial surface. Overall, these boundary conditions allow rotation at the knee and bending of the tibia during axial compression. For validation purposes, the measured longitudinal strains were compared to the finite element surface strains over the same surface area occupied by the gages (0.38mm × 0.5mm). In addition, the strains (longitudinal, minimum and maximum principal) and von Mises stresses were determined at four locations around the defect (Figure 2a, white stars) and on the medial periosteal surface on the opposing cortex (Figure 2a, red star).

**Figure 2.**
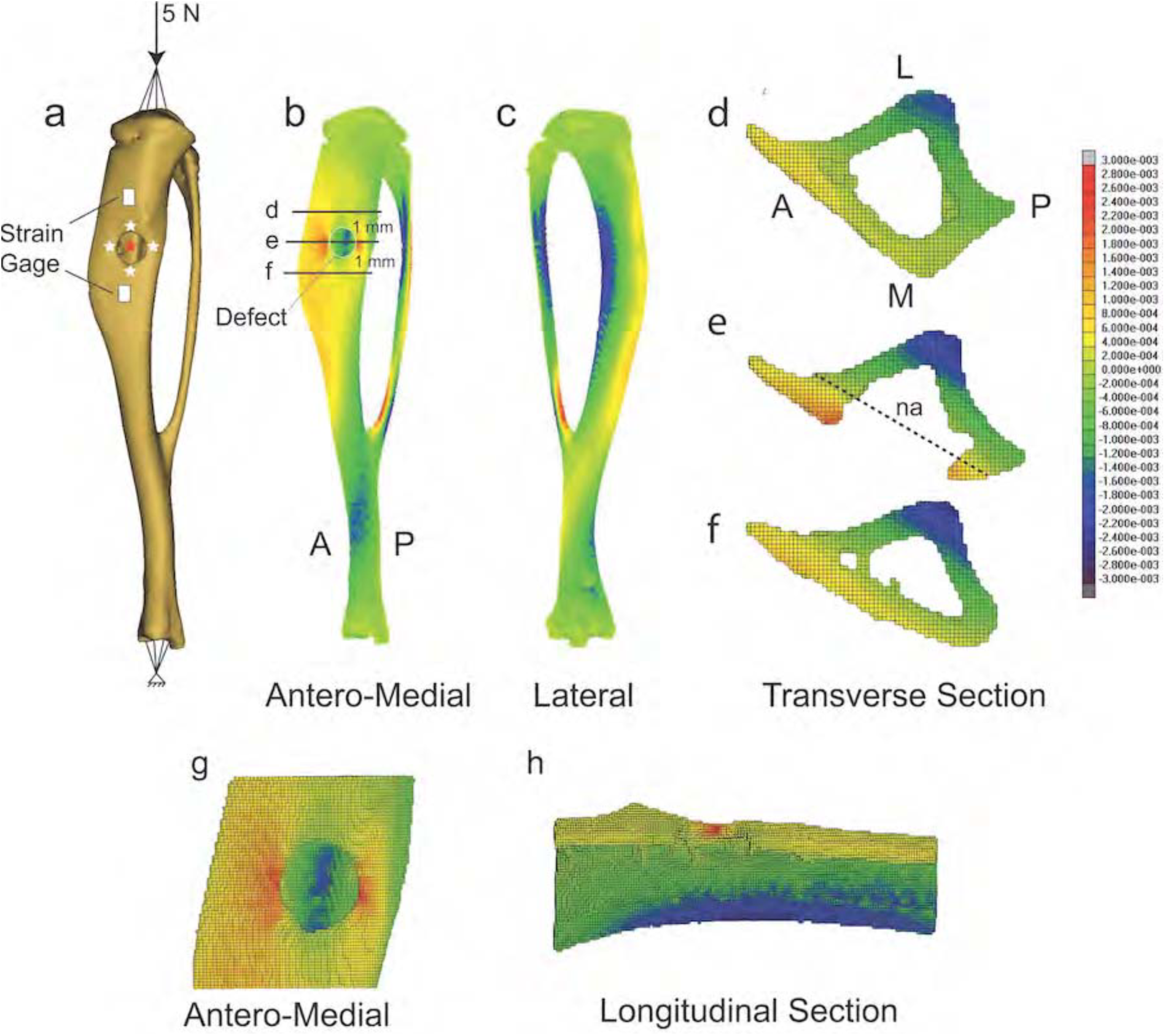
Distribution of longitudinal strain in a representative tibia containing a monocortical defect and in sections containing, or near, the defect. (a) Rendered tibia showing defect, applied compressive 5 N peak load, strain gage placement, and positions along the defect border (white stars), and opposing cortex (red star), where strain components were evaluated. (b) Antero-medial view. (c) Lateral view. Transverse section located: (d) 1 mm proximal to the center of the defect, (e) at the defect center, and (f) 1 mm distal to the defect. (g) Magnified antero-medial view of the bone segment containing the defect. (h) Longitudinal section of segment containing the defect. Red indicates tensile strain, blue indicates compressive strain. na = neutral axis.

### 2.4 In vivo tibial loading

Tibiae were subjected to four consecutive days of *in vivo* axial compressive cyclic loading (5 N peak load, 2 Hz, 60 cycles) via load feedback using an electromagnetic mechanical loading device (TA EnduraTEC 3200, Enduratec Systems Group). The tibia was secured at the flexed knee and ankle joints, and an axial load was applied through the knee creating bending and compressive strains throughout the tibia [16,17]. Loading began on PSD 2, 5, or 10, each of which corresponds to distinct stages of repair: (1) PSD 2 to 5, hematoma formation and inflammatory exudation; (2) PSD 5 to 8, bone matrix deposition; and (3) PSD 10 to 13, matrix mineralization.

### 2.5 In vivo longitudinal microCT

MicroCT scanning and analyses were performed in concordance with published guidelines [18].Transverse high-resolution microCT images of bilateral defects were simultaneously acquired (210 slices, 55 kVp, 300 ms integration time, 10.5 μm isotropic voxel size) on PSD 2, 5, 8, 10, and 14 using an *in vivo* microCT scanner (vivaCT 40, Scanco USA, Inc.). The time from the onset of anesthesia to recovery was approximately 25 minutes, and the total scanning time was approximately 15 minutes. The region of interest (ROI) was centered at mid-defect, and included the defect, regenerate (localized irregular tissue within the defect consisting of woven bone matrix, connective tissue, and cells), and surrounding cortices (Figure S1) selected through segmentation using a constant global threshold based on the manufacturer’s recommendations and preliminary studies comparing the original and segmented scan side-by-side to ensure segmentation accuracy. Percent change in bone volume relative to PSD 2 was calculated for each defect at PSD 5, 10 and 14.

### 2.6 Tissue processing and histomorphometry

At PSD 2, 5, 10, and 14, tibiae were harvested and fixed in 4% paraformaldehyde overnight and decalcified in 19% EDTA for 14 days at room temperature. Fixed bones were processed for paraffin embedding and sectioned in the longitudinal direction into 6 μm serial sections. Consecutive sections at mid-defect were stained for: (1) aniline blue (every 9^th^ slide) to evaluate new bone formation, (2) Movat’s pentachrome staining to visualize gross tissue appearance, (3) proliferating cell nuclear antigen (PCNA) to assess cellular proliferation, (4) Runx2 to assess osteogenic differentiation, (5) alkaline phosphatase (ALP) to examine osteoblast activity, (6) tartrate-resistant acid phosphatase (TRAP) to evaluate osteoclastic activity, and (7) lectin (DyLight 594 Ulex Europaeus Agglutinin) to assess vascularization of the defect. Sections were imaged using a Leica DM5500 microscope at 5x, and images were evaluated using ImageJ. Formation of bone and cartilage within the tibial defects was assessed from aniline blue and pentachrome stained sections, respectively, by counting selected pixels as previously described [15]. Gross histological appearance of PCNA, Runx2, ALP, and TRAP staining, and quantified bone and cartilage area were compared between experimental groups.

### 2.7 Data analyses

Differences in (1) percent change in bone volume, (2) bone area, and (3) cartilage area on PSD 5, 10 and 14 were compared by a one-way ANOVA with loading group (control, PSD 2 to 5, PSD 5 to 8, and PSD 10 to 13) as the main factor (Prism 6 Statistical Software, GraphPad Software, Inc.). Data are presented as mean +/‐ standard deviation. Statistical significance was assumed for *p* < 0.05.

## 3.0 Results

### 3.1 FE model validation and strain distribution in cortical bone surrounding the defect

To better understand the mechanical environment of a monocortical tibial defect subjected to exogenous loading, we developed FE models of three separate tibiae each containing a single defect. Longitudinal tensile strains measured on the cortical surface (Figure 2a) in response to a 5 N peak load applied in axial compression through the knee joint were lower for the proximal gage (326±34 με) compared to the distal gage (598±312 με). Corresponding FE results for the proximal (311±83 με) and distal (608±155 με) gage locations corresponded with the measured experimental strain.

Bending at the proximal tibia was evident with tensile and compressive strains on the medial and lateral surfaces, respectively (Figure 2b, c). The neutral axis was parallel with the antero-medial aspect of the tibia, and shifted from the center of mass of the section (Figure 2e) indicative of a combination of compression and bending. Near the defect, there was a tensile strain concentration along the anterior and posterior borders (~2200 με), while the strain on the proximal and distal borders were much smaller (~50-100 με) (Table 1). Strain on the periosteal cortex opposing the defect had the highest strain magnitudes (~-2600 με) and were compressive in nature. The concordance between longitudinal strain and maximum principal strain on the medial side, and longitudinal strain and minimum principal strain on the lateral side, demonstrated that the proximal tibia is loaded in compression and bending. In sum, loading produces a strain field around the defect that is high on the anterior and posterior borders and low on the proximal and distal borders (Figure 2g, h).

**Table 1.** Bone strain components at four positions^a^ around the monocortical tibial defect edge and on the opposing cortex at the posterior aspect of the tibia.

| | $\epsilon_{zz}^b$ ($\mu\epsilon$) | $\epsilon_{min}^c$ ($\mu\epsilon$) | $\epsilon_{max}^d$ ($\mu\epsilon$) | von Mises Stress (MPa) |
| --- | --- | --- | --- | --- |
| **Anterior** | $2,353 \pm 689$ | $-623 \pm 112$ | $2,140 \pm 450$ | $37.3 \pm 9.7$ |
| **Posterior** | $2,033 \pm 503$ | $-540 \pm 239$ | $2,260 \pm 589$ | $35.9 \pm 8.0$ |
| **Proximal** | $93 \pm 77$ | $-233 \pm 188$ | $155 \pm 21$ | $6.2 \pm 1.7$ |
| **Distal** | $44 \pm 51$ | $-270 \pm 61$ | $50 \pm 29$ | $7.5 \pm 0.9$ |
| **Opposing Cortex** | $-2,543 \pm 450$ | $-2,657 \pm 450$ | $690 \pm 69$ | $42.9 \pm 5.5$ |

<sup>a</sup>See Figure 2 for position information
<sup>b</sup>Strain ( $\epsilon$ ) in the longitudinal direction (zz) and parallel to the long axis of the tibia in units of microstrain ( $\mu\epsilon$ )
<sup>c</sup>Minimum principal strain
<sup>d</sup>Maximum principal strain

### 3.2 Sample-specific changes in bone volume during healing

To evaluate longitudinal changes in bone volume at the defect site, *in vivo* longitudinal microCT scanning of defects was performed at PSD 2, 5, 8, 10 and 14. Significant differences in percent change bone volume between experimental groups at each time point were not detected, and high variability within groups was observed (Figure 3).

**Figure 3.**
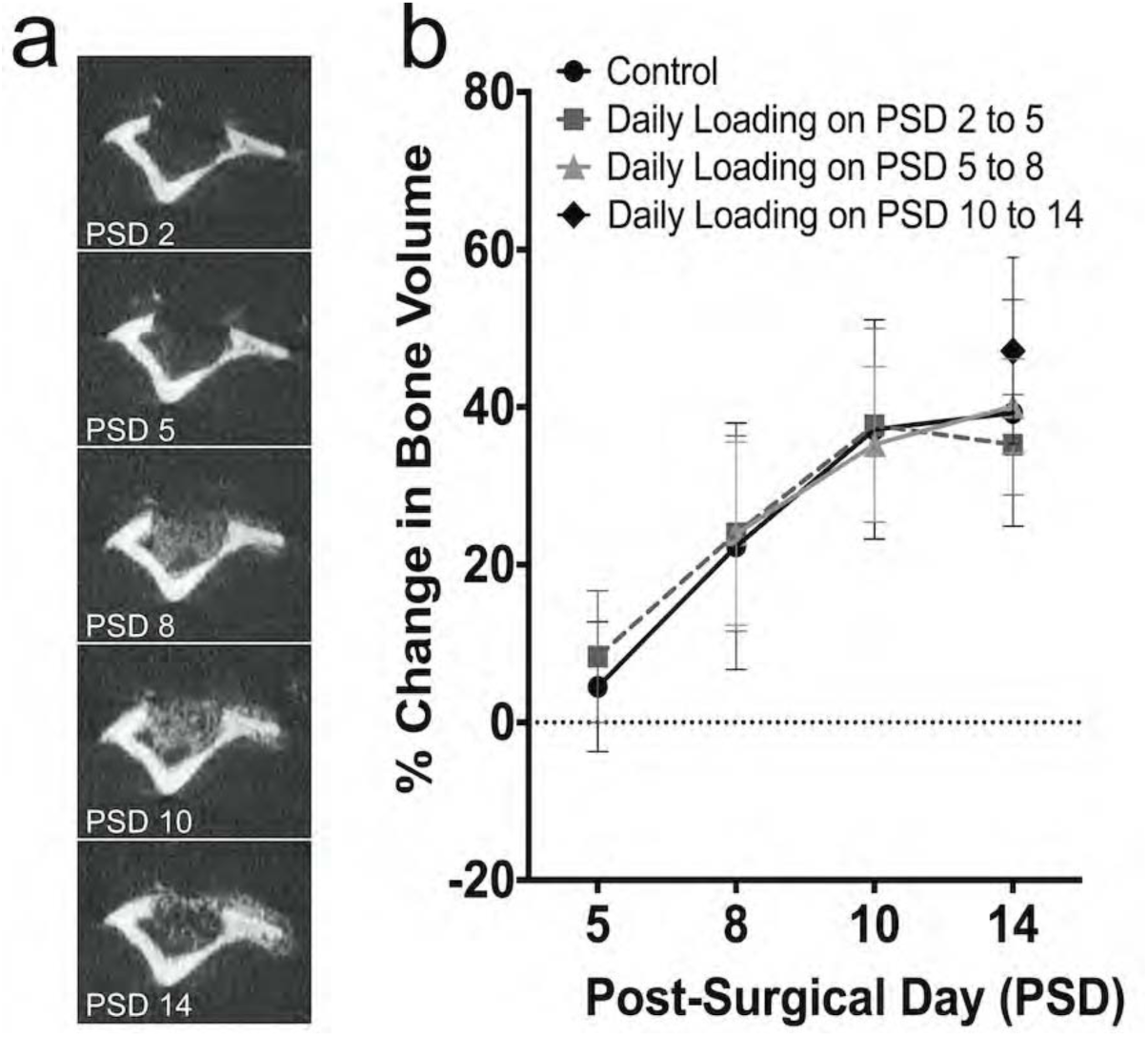
Percent change in bone volume in control and loaded tibiae during the course of healing by longitudinal microCT scanning. (a) Longitudinal microCT scans at PSD 2, 5, 8, 10 and 14 in a non-loaded control tibia show an increasingly mineralized regenerate at the defect site. (b) Percent change in bone volume relative to PSD 2 for all experimental groups increased at PSD 5, 8 and 10, and leveled-off by day 14. Data represent mean ± SD (*n*=8 control, *n*=16 loaded on PSD 2 to 5, *n*=10 loaded on PSD 5 to 8, *n*=9 loaded on PSD 10 to 13).

### 3.1 Daily loading during the inflammatory phase (PSD 2 to 5) delays hematoma clearance and bone matrix deposition, stimulates cellular proliferation and osteoclast activity, and promotes cartilage formation.

To gain insight into the effects of loading during the early stages of defect repair, defects were subjected to loading on PSD 2 to 5, when hematoma formation, inflammatory exudation, and granulation tissue formation are underway. At PSD 2, control defects (Figure 4, Figure S2) were filled with a hematoma (Figure 4a) with robust cellular proliferation within the hematoma (Figure 4e) and at the periosteal surface (Figure 4i). with relatively low levels of osteogenic differentiation (Figure 4m) and osteoclast activity (Figure 4q) detected. At PSD 5, control defects showed bridging with woven bone and new bone formation at the periosteal surface (Figure 4b, j) with cellular proliferation evident within the defect (Figure 4 f, j). Osteogenic differentiation (Figure 4n) peaked within the defect at PSD 5 and small numbers of TRAP+ cells were evident (Figure 4r) within the defect suggesting the beginning of osteoclastic matrix remodeling. At PSD 10, an enlarged regenerate was present (Figure 4c) with osteoblast (ALP) and osteoclast (TRAP) activity underway (Figure 4 o, s). At PSD 14, the regenerate was substantially remodeled (Figure 4d) cellular proliferation detected in soft tissue above the periosteal surface (Figure 4 l). Remodeling of the regenerate was still underway (Figure 4p, t).

**Figure 4.**
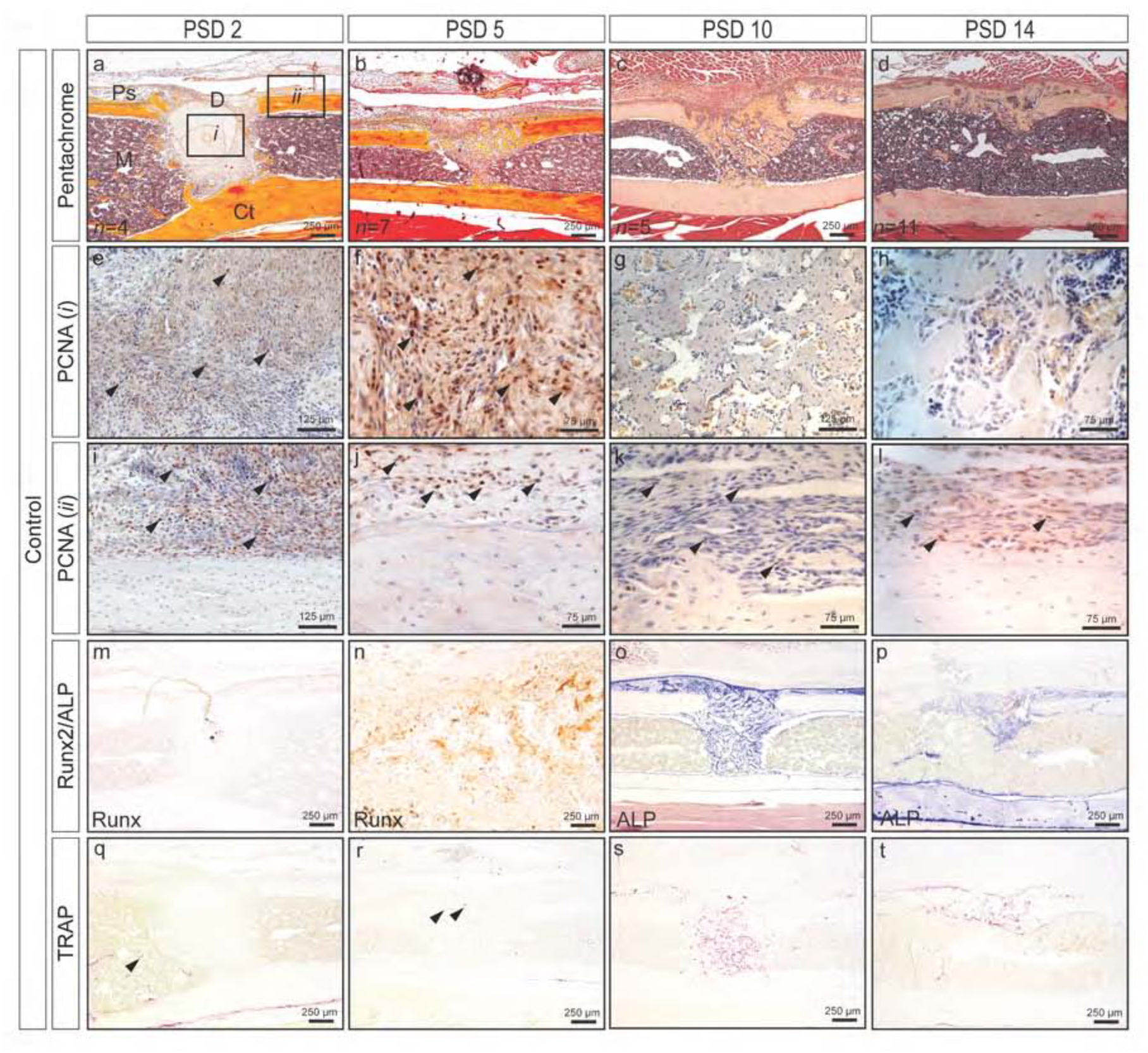
Time course of monocortical defect healing in control mouse tibia, (a-d) Pentachrome staining reveals defect healing through intramembranous bone formation. (e-l) Robust cellular proliferation (black arrows, PCNA+ cells are brown) within the defect (*i*) and at the periosteal surface (*ii*) up to PSD 5. Cellular proliferation in fibrous tissue above the periosteum (k, l), but not within the defect (g, h) at PSD 10 and 14. (m-p) Osteogenic differentiation (Runx2) (m, n) and osteoblast activity (ALP) (o, p). (q-t) Low levels of osteoclast activity are evident within the marrow cavity and defect at PSD 2 and 5 (black arrows), and increases at PSD 10 and 14 as the regenerate begins to remodel. Quantified bone and cartilage data for controls are presented in Figures 6 thru 8 alongside data for loading groups. D = defect; M = marrow; Ps = periosteal surface; Ct = cortical bone.

Defects subjected to daily loading from PSD 2 to 5 (Figure 5) exhibited a hematoma within the defect (Figure 5a, d) and an elevated periosteum consisting of proliferating cells (Figure 5g) at PSD 5. The hematoma likely resulted from either newly perforated blood vessels in response to loading and/or a delay in removal of the primary hematoma formed immediately after surgery. Proliferating cells were observed within the defect at all time points post-loading and within the elevated periosteum and surrounding cartilage nodules suggesting that loading activated proliferation even when strains were relatively low (50-100 με). Relative to controls at PSD 5, less osteogenic differentiation within the defect and more TRAP staining at the periosteal and endosteal envelopes were observed. Cartilage and fibrocartilage formation was observed above the defect and periosteal surface at PSD 10 (Figure 5b, p) indicating that loading during the inflammatory phase pushes osteochondroprogenitors into a cartilage phenotype. Finally, robust remodeling as assessed by ALP and TRAP staining was observed at PSD 14. Overall, mechanical loading during the early stages of repair appears to disrupt the injury site resulting in a prolonged healing program.

**Figure 5.**
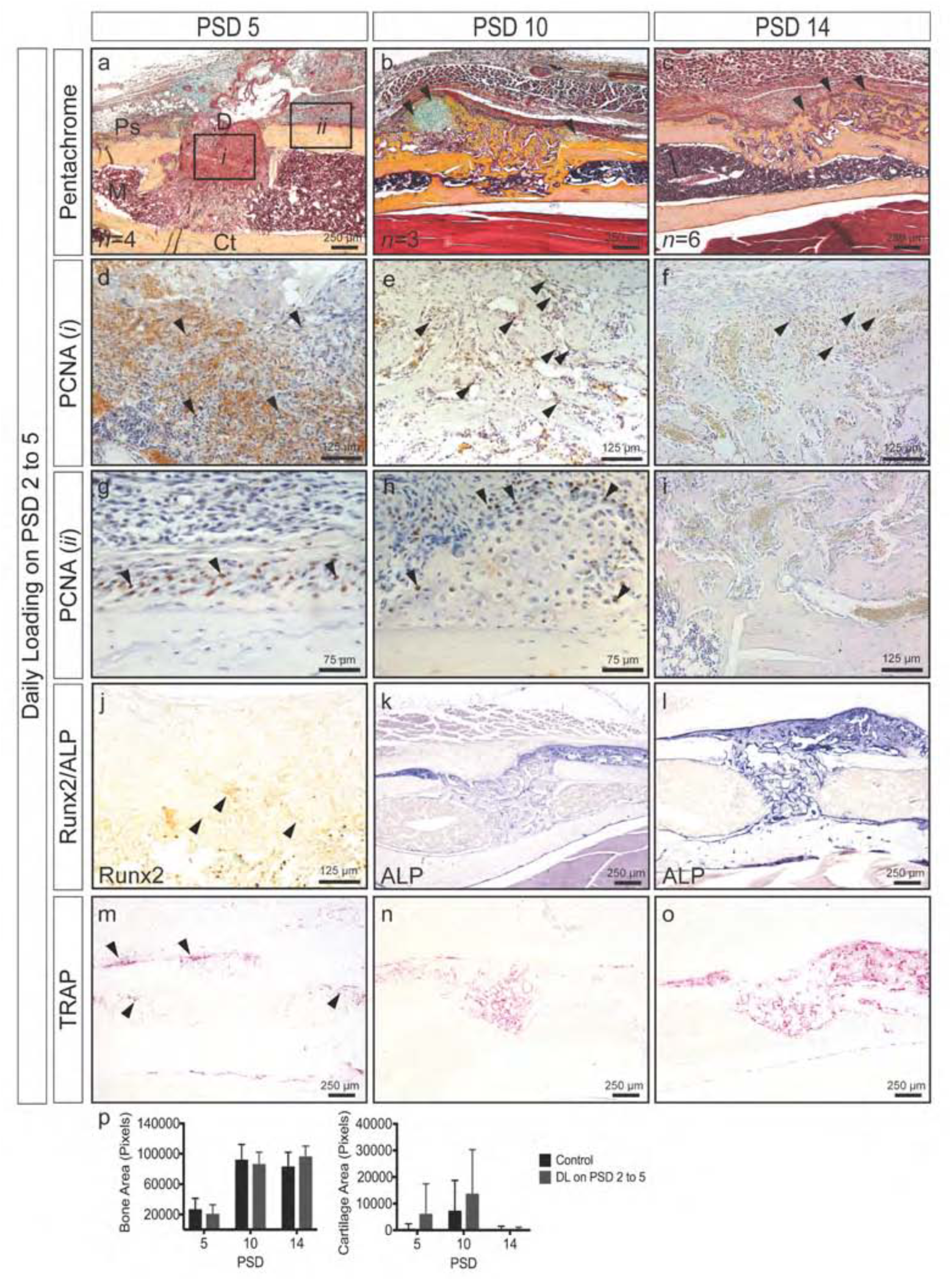
Time course of monocortical defect healing in response to daily loading at PSD 2 to 5. (a-c) Pentachrome staining reveals robust cartilage formation (black arrows) at PSD 10 and the continued presence of woven bone (black arrows) at PSD 14. (d-i) Robust cellular proliferation (black arrows, PCNA+ cells are brown) within the defect (*i*) at all time points. An elevated periosteum (*ii*) containing proliferating cells (black arrows) at PSD 5. Proliferating cells (black arrows) at the cartilage periphery above the periosteal surface (h). (j-l) Osteogenic differentiation (Runx2, black arrows) and activity (ALP). (m-o) Enhanced osteoclast activity at PSD 5 (black arrows). (p) Quantified bone and cartilage area. Data represent mean ± SD. Differences between groups at each time point tested by one-way ANOVA. Significance accepted at α = 0.05. D = defect; M = marrow; Ps = periosteal surface; Ct = cortical bone.

### 3.2 Daily loading during the matrix deposition phase (PSD 5 to 8) stimulates cellular proliferation, promotes cartilage formation, and enhances bone formation.

To better understand the effects of mechanical loading during the matrix deposition phase, defects were subjected to loading on PSD 5 to 8. Histological evaluation (Figure 6) showed cartilage and fibrocartilage formation at or near the periosteal surface at PSD 10 (Figure 6a) and 14 (Figure 6b), though differences between groups did not reach significance (k). Exuberant cellular proliferation was observed within the defect (Figure 6c, d) and at the periphery of cartilage nodules (Figure 6e, f) above the periosteum. Loaded defects displayed significantly greater bone area (*p* = 0.02) at PSD 10 (k). This result together with enhanced proliferation within the defect suggests that loading activates proliferation. Remodeling, as assessed by osteoblast (Figure 6g, h) and osteoclast activity (Figure 6i, j), was evident throughout the regenerate and at the periosteal and endosteal surfaces at PSD 10 and 14. Thus, mechanical loading during matrix deposition activates additional bone formation and modest levels of cartilage formation.

**Figure 6.**
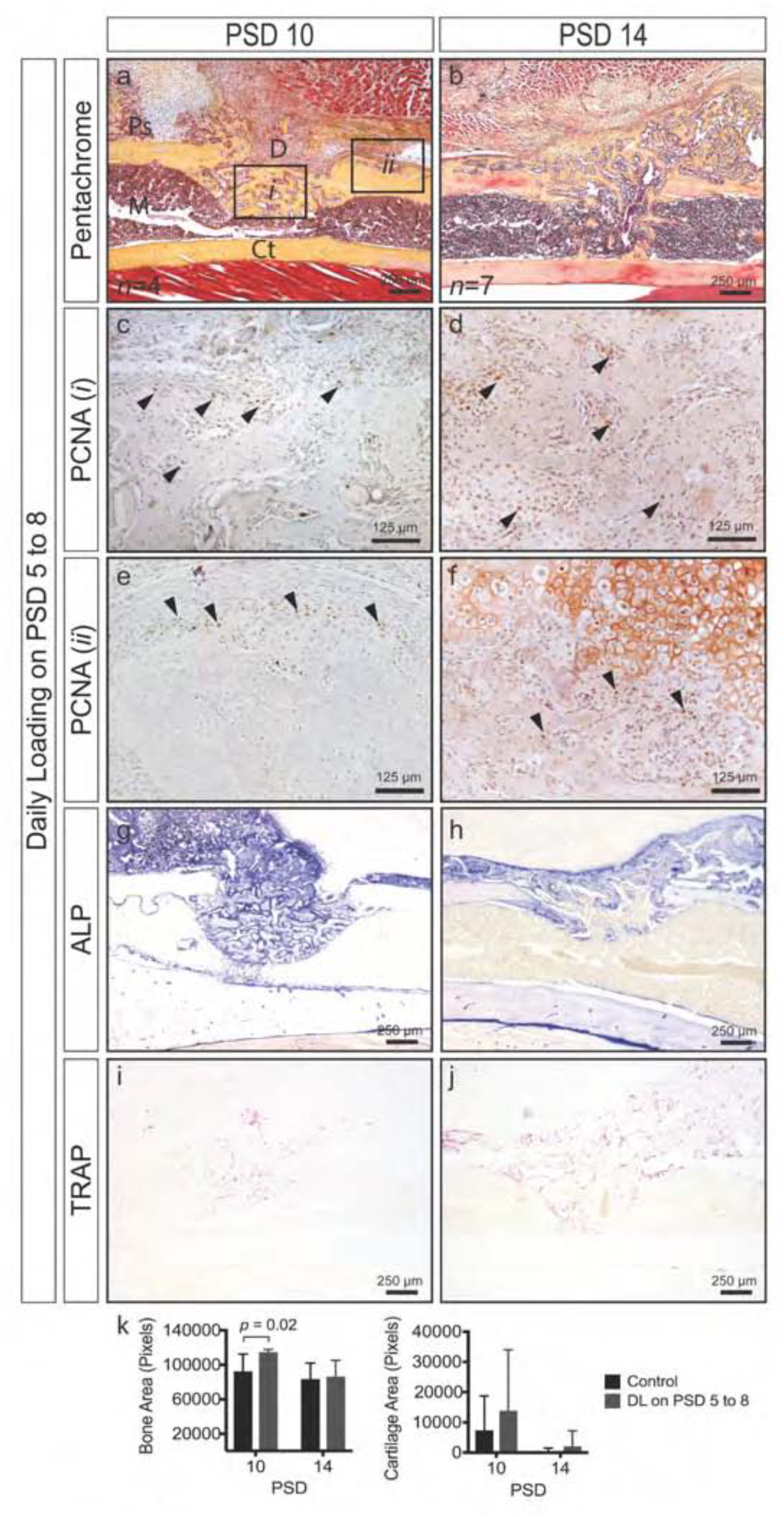
Time course of monocortical defect healing in response to daily loading at PSD 5 to 8. (a, b) Pentachrome staining reveals cartilage formation (black arrows) at PSD 10 and the continued presence of woven bone and fibrocartilage (black arrows) at PSD 14. (c-f) Robust cellular proliferation (black arrows, PCNA+ cells are brown) within the defect (*i*) and above the periosteal surface near cartilage nodules (*ii*). (g-j) Osteoblast (ALP) and osteoclast (TRAP) activity throughout the regenerate. (k) Quantified bone and cartilage area; significantly greater bone area (*p* = 0.02) in loaded versus control defects at PSD 10. Data represent mean ± SD. Differences between groups at each time point tested by one-way ANOVA. Significance accepted at α = 0.05. D = defect; M = marrow; Ps = periosteal surface; Ct = cortical bone.

### 3.3 Daily loading during the remodeling phase (PSD 10 to 13) stimulates cellular proliferation and prolongs the remodeling phase.

To examine the effects of mechanical loading during the remodeling phase of repair, defects were subjected to loading on PSD 10 to 13 when the regenerate is undergoing rapid turnover. Relative to controls (Figure 3), loaded defects showed abundant amounts of woven bone within the defect and at the periosteal surface (Figure 7a), with proliferating cells dispersed throughout the regenerate (Figure 7b, c). Much of the regenerate was undergoing osteoblast‐ and osteoclast-driven remodeling as assessed by ALP and TRAP staining. Though abundant amounts of woven bone was present, cartilage formation was not observed at PSD 14 suggesting that the regenerate had reached a point of maturation at which fewer osteochondroprogenitors were present and the stiffening matrix reduced overall load-induced strain.

**Figure 7.**
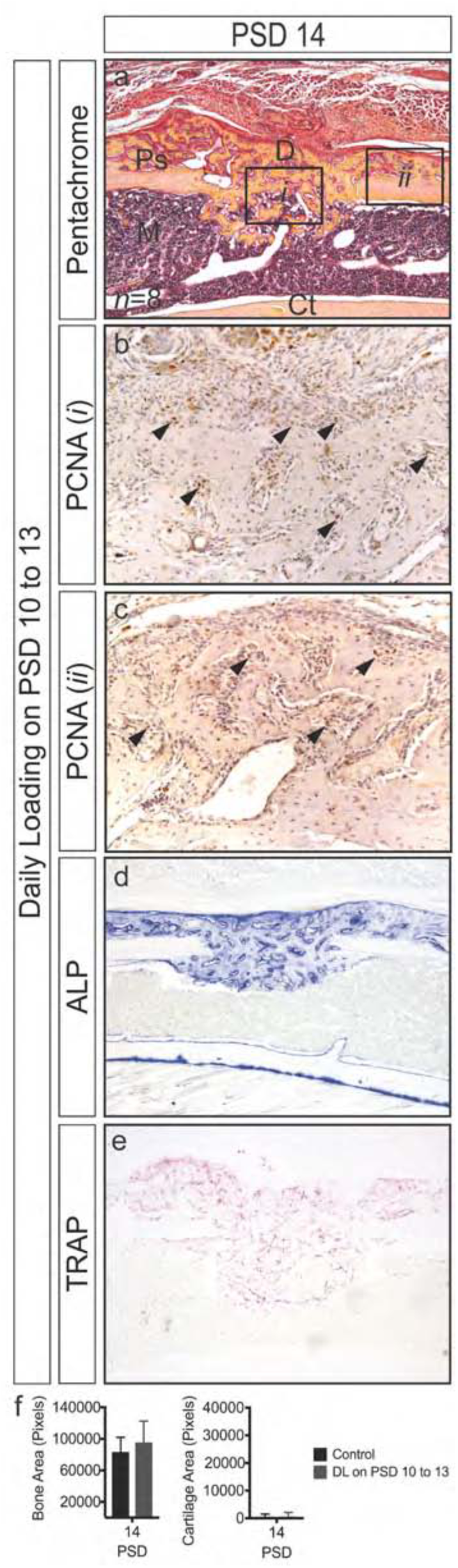
Monocortical defect healing in response to daily loading at PSD 10 to 13. (a) Woven bone within the defect and at the periosteal surface at PSD 14 (b) Robust cellular proliferation (black arrows) within the defect (*i*) and in regions of woven bone above the periosteal surface (*ii*). (d, e) Osteoblast (ALP) and osteoclast (TRAP) activity are apparent throughout the regenerate. (f) Bone area is similar between control and loaded defects. Only small regions of cartilage formed in response to loading. Data represent mean ± SD. Differences between groups at each time point tested by one-way ANOVA. Significance accepted at α = 0.05. D = defect; M = marrow; Ps = periosteal surface; Ct = cortical bone.

## 4.0 Discussion

Compressive strain, tensile strain, hydrostatic pressure, shear strain, and fluid dynamics have all been implicated as important mechanical cues regulating bone regeneration [2,6,19], and there is a rapidly growing body of quantitative work describing relationships between mechanical factors, tissue formation, and tissue-specific differentiation during bone healing [7–10,12,13,20]. Overall, these studies have suggested that low stress and strain lead to direct intramembranous bone formation, compressive stress and strain lead to chondrogenesis, and high tensile strain leads to fibrous tissue formation [6,9]. Still, our understanding of the effects of mechanical loading on repair is incomplete, particularly when applied at different stages of healing. To address this question, we used a mouse tibial loading model coupled with a monocortical defect, which revealed that loading during the earliest stages (PSD 2 to 5) of repair disturbs the injury site as evidenced by a persistent hematoma. Our analyses could not distinguish whether the persistent hematoma was because of a delay in clearance of the primary hematoma, or because the mechanical loading ruptured the newly forming microvasculature, which resulted in a second hematoma. We also found that loading activates a strong periosteal response near the defect rim resulting in enhanced cellular proliferation and cartilage formation, even at relatively low tensile strain (50-100 με). Cartilage and fibrocartilage formation ultimately delays healing of the defect as evidenced by persistence of a large woven bone regenerate two weeks after surgery when control regenerates have already undergone substantial remodeling. These data are consistent with other studies reporting that loading immediately after fracture inhibits vascular invasion and bone formation [10], and activates greater cartilage formation [12]. Whether vascular disruption or the presence of cartilage inhibits bone formation in unclear, but both hypoxia [21] and cartilage-specific anti-angiogenic factors [22–24].

Loading during the matrix deposition stage (PSD 5 to 8), when the regenerate has bridged the defect, leads to a significant increase in bone formation by PSD 10, a result consistent with reports showing that loading during osteogenesis enhances bone formation [11]. A concomitant remodeling of the vasculature and larger vessels was also reported, which may be interpreted to mean that a more mature vascular bed and larger vessels allows for greater ingress of progenitors, thereby enhancing tissue formation and healing. Enhanced cellular proliferation and cartilage formation was also observed with this loading regimen suggesting that resident osteochondroprogenitors, even at this later stage of healing, can undergo proliferation and/or chondrogenic differentiation. Loading during the remodeling phase (PSD 10 to 14) lead to increased cellular proliferation throughout the regenerate, but not cartilage formation, suggesting that osteochondroprogenitors may have been fewer in number and that the stiffer matrix resisted deformation. Additional work is needed to address the degree of stiffening of the regenerate as healing progresses.

FEA showed that a monocortical defect in the mouse tibia acts as a stress concentrator resulting in high strains at the anterior and posterior defect edges and low strains at the proximal and distal defect edges. Histological assessment showed that defect-adjacent periosteal surfaces experiencing ~50-100 με coincided with areas of distinct and exuberant periosteal chondrogenic differentiation in a model that typically heals through intramembranous bone formation [14]. These strain values fall at the low end of a range of values (300 to 24,000 με) reported to enhance cartilage formation at an injury site [10–13,25,26]. Inflammation is an important regulator of stem cell function [27], and the proximity of the affected periosteal sheath to the defect and inflammatory environment may have sensitized periosteal resident progenitors to mechanical stimulation by altering gene expression and cell behavior. Additional work is required to understand the relationship between pro-inflammatory cytokines and stemness in the context of fracture repair and loading.

Even though a significant increase in bone volume was detected by aniline blue staining in bones loaded at PSD 5 to 8 relative to controls, this difference was not observed with longitudinal microCT analysis. One possible explanation is the heterogeneous nature of the regenerate with regard to structure and varying degrees of mineralization, and the challenge it presents with regard to thresholding and partial volume effects. In this sense, histological assessment provides important validation of microCT outcomes, and both should be presented when possible. Movement artifacts may also have contributed to data variability.

A variety of *in vivo* models and computational analyses have been implemented to study mechanobiology of bone adaptation [16,17,28–31] and repair [6,8,10,12,25]. Of these, murine bone injury models are particularly attractive given their relatively low cost and the ability to genetically engineer transgenic and targeted knockout mouse lines, allowing study of molecular mechanisms governing bone repair. In this study we utilized an *in vivo* murine tibial loading model that is well characterized and widely used to study mechanical adaptation of bone [16,28,31,32]. In this regard, the model easily accommodates a variety of user-defined loading regimens that can be applied during distinct stages of defect healing. One key advantage of this model is the ability to correlate spatial strain distribution in response to loading with distinct tissue formation in the context of repair. Furthermore, the effects of a wide range of variables (load magnitude, cycle number, frequency, and latency period) on repair are testable using this model. Finally, its technical simplicity (reproducibility of the defect, does not require stabilization, permits normal ambulation immediately after surgery, confers minimal risk of failure post-surgery, is void of radio-opaque hardware permitting *in vivo* longitudinal skeletal imaging) makes it highly accessible. This model allows the study of distinct bone envelopes in response to controlled mechanical loading in an injury setting. In addition, the inhomogeneity in strain due to the asymmetry of the bone [30] will be advantageous for evaluating a range of load-induced stress and strain values on bone repair.

Limitations with the presented model are as a stable cortical bone repair model, the complexity of a full fracture environment is not realized; however, we believe this limitation is outweighed by the ability to apply controlled and highly reproducible loading protocols as well as simplifying development of FE models given the continuous cortical shell. A second limitation is that the CT-based FE analysis does not incorporate the contribution of the regenerate or bone marrow to the overall mechanical environment, particularly at later stages of healing when the regenerate begins to stiffen. These aspects of the model will be addressed in future work.

In summary, our findings suggest that early loading disrupts the injury site and pushes periosteal-resident osteochondroprogenitors into a cartilage phenotype while loading during the bone matrix formation phase pushes progenitors into a bone phenotype. Due to its technical simplicity and spatially confined repair process, the described mechanobiological model may offer advantages for studying distinct bone envelopes in response to precisely controlled mechanical loading regimens in the absence of hardware fixation.

## Acknowledgements

This work was supported by the AO Foundation (Switzerland) Award S-13-57C (ABC) and the Department of Veterans Affairs, Veterans Health Administration, Office of Research and Development, Career Development Award-2 A6842W (ABC).

## Author Contributions

Study design: CL, ABC. Data collection: CL, VF, RC, LK, GB, DH, BL, GS, PL, KAM, ABC. Data analysis and interpretation: CL, VF, PL, KAM, JAH, ABC. Drafting the article: CL, KAM, ABC. Critical revision of the article: CL, PL, KAM, JAH, ABC. All authors approved the final version of the article. Correspondence should be addressed to ABC.

